# Topological Analysis of the Human Lymph Node Reticular Network Predicts Outcome in Breast Cancer

**DOI:** 10.1101/2025.07.07.663481

**Authors:** Amy Llewellyn, Sharon D’Costa, Ringo Lam, Jasmine Gore, Veronika Lachina, Daniel Shewring, Sophie E. Acton, Kalnisha Naidoo

## Abstract

Axillary lymph nodes (ALN) initiate local immune responses in breast cancer (BC) but how and when ALN become dysfunctional, facilitating metastasis, is unclear. We use unbiased computational approaches to quantify features of ALN stromal architecture. We identify PDGFRβ as a robust, immunomarker for human fibroblastic reticular cells (FRC) and use it to quantify how FRC network topology changes during BC progression and after treatment. ALN (n = 331) from 179 BC patients and 23 benign reactive controls were assessed for FRC network metrics, including lacunarity and branchpoints, alongside de-identified clinico-pathological data. We find in node-negative, triple-negative BC, neoadjuvant treatment induced denser FRC networks which correlated with improved survival. Conversely, denser FRC networks in node-positive patients correlated with worsened survival, regardless of BC subtype or treatment. Further, increased FRC alignment within metastases improved survival. We show that FRC network topology predicts prognosis in BC, providing a new avenue for mechanistic, translational research.

## INTRODUCTION

In breast cancer (BC), axillary lymph node (ALN) status is central to guiding management decisions since sentinel lymph node (SLN) involvement is the most significant prognostic factor in early-stage disease. However, while ALN are the first site of BC metastasis, they are also highly organized, tightly regulated secondary lymphoid organs capable of generating powerful adaptive immune responses against tumor cells. Furthermore, ALN modulate fluid homeostasis in the arm and breast. Consequently, surgical clearance of the axilla can result in lymphoedema of the arm but the effects on immune function are still unclear ^1^.

Over the past 30 years, BC treatment has evolved significantly, now integrating tailored combinations of neoadjuvant and adjuvant systemic therapies based on disease stage and hormone receptor status (estrogen receptor positive (ER+), human epidermal growth factor receptor 2 positive (HER2) and triple negative breast cancer (TNBC)). In parallel, management of the axilla has also changed, with the number of ALN clearances (ALNC) decreasing in favor of less invasive techniques such as SLN biopsy (SLNB) and, more recently, targeted axillary dissection ^2^. Clinical trials, such as Z0011 and AMAROS, have led to this paradigm shift by challenging the necessity of ALNC in patients with minimal axillary involvement ^3,4^. In particular, the fact that isolated tumor cells (ITCs; < 200 cells or < 0.2 mm in extent) and micrometastasis (0.2–2 mm in extent) do not impact recurrence or overall disease-free survival, if appropriate systemic therapies are given, supports conserving the axilla ^5^. Despite this however, precise histopathological examination of every surgically removed ALN is still mandated to ensure accurate prognostication and staging. Furthermore, with neoadjuvant immunotherapy now being standard of care for TNBC, the potential of ALN microenvironment to either activate or suppress the immune response is clinically relevant ^6^. However, our biological understanding of the mechanisms of ALN metastasis, and the factors that determine if a patient will develop an effective anti-tumor immune response or not, remains poorly understood.

Fibroblastic reticular cells (FRC) are integral to ALN structure and function. These specialized stromal cells secrete and enwrap extracellular matrix to form a network of reticular fibers that facilitate efficient immune cell localization and antigen transport (reviewed in ^7^). Lymph enters the lymph node via the subcapsular sinus (SCS), where sinus-lining cells act as a molecular sieve, restricting molecules larger than ∼70 kDa from accessing the conduit system ^8–10^. Smaller soluble molecules filter into the FRC ensheathed conduit network and flow in a unidirectional, controlled manner through the lymph node parenchyma to high endothelial venules (HEV). In addition to their structural scaffolding role, FRC orchestrate immune cell interactions through producing chemokines such as chemokine ligand (CCL)19 and CCL21 which regulate lymphocyte migration. Single cell RNA sequencing (scRNAseq) has elucidated distinct subsets of murine FRC, each with highly specific roles in maintaining immunological niches ^11^. However, the canonical marker of murine FRC remains Podoplanin (Pdpn). In mice, this membranous glycoprotein has been shown to have central role in the maintenance and dynamic remodeling of the conduit network, and in immunoregulatory properties of FRC ^12–14^.

In contrast, our knowledge of human FRC biology and their response to immune activation and cancer is still evolving. Studying human FRC is challenging due to the ethical constraints on accessing lymphoid tissue. In BC patients, every excised ALN must be formalin-fixed and paraffin-wax embedded (FFPE) and examined in its entirety for accurate pathological staging ^15^. This limits the number of patient-derived ALN samples available for research. In addition, FRC make up only 1-5% of human lymph nodes and are significantly more difficult to isolate from primary tissue than immune cells ^16,17^. Furthermore, it is widely recognized that subclassifying human fibroblasts is challenging due to the lack of unique, robust markers (reviewed in ^18^). Attempts to isolate human FRC have traditionally used PDPN to define the population, but PDPN expression is low in human FRC ^19–21^. However, very recent scRNAseq of reactive human lymph nodes has revealed that platelet-derived growth factor receptor β (PDGFRβ) is expressed by all subsets of lymph node fibroblasts ^22^. Finally, the collagenous reticular network formed by murine FRC also exists in human lymph nodes, yet our understanding of how its structure and composition change in disease remains limited. Histopathologists have historically used tinctorial stains such as reticulin to visualize this delicate network ^23^ and Masson’s trichrome to highlight mature fibrosis ^24^, but these methods have not been correlated with functional and/or clinical data, or FRC topography.

There are limited data on freshly resected ALN containing BC metastases, which identify four fibroblast subsets with prognostic significance based on the differential expression of five fibroblast markers: fibroblast activation protein α1 (FAP), integrin β1, smooth muscle actin (SMA), PDGFRβ, and PDPN ^25^. However, this study was limited by a small sample size, the absence of benign reactive controls and did not define the spatial distribution of these FRC populations within ALN.

Herein, we define robust immunohistochemical (IHC) markers of human FRC and characterize their expression patterns in ALN from a large cohort of BC patients, as well as reactive, control, tumor-free lymph nodes from patients with benign disease. We comprehensively map and quantify the effect of BC and neoadjuvant chemotherapy (NACT) on FRC network topology in both involved and uninvolved ALN, linking changes in FRC network topology to patient survival.

## RESULTS

### Optimized IHC Markers of Human FRC Subsets

We analyzed 331 FFPE ALN samples from 179 BC patients diagnosed and treated at King’s College Hospital (KCH), encompassing all three molecular subtypes, as well as different TNM stages and variable responses to NACT (**Table 1**). No patients had received checkpoint inhibitor therapy (CPI). 23 tumor-free lymph nodes from patients with benign disease, who had never had cancer, served as reactive controls. Given the paucity of robust IHC stains for human lymph node FRC, we optimized protocols for five previously reported FRC markers ^26^ and compared staining patterns across serial sections to reticulin, a classic tinctorial stain which is known to highlight the collagen backbone of lymphoreticular organs ^23^.

**Table 1.** Clinico-pathological Characteristics of BC Patient Cohort.

| Clinicopathological Characteristic |  | No. (%) |  |
| --- | --- | --- | --- |
| **Age** |  |  |  |
| < 50 |  | 44 (25) |  |
| > 50 |  | 135 (75) |  |
| **Pathological T stage** |  |  |  |
| ypT0 |  | 55 (31) |  |
| pT1 | ypT1 | 42 (23) | 17 (9) |
| pT2 | ypT2 | 44 (25) | 8 (4) |
| pT3 | ypT3 | 6 (3) | 6 (3) |
| pT4 | ypT4 | 1 (1) | 0 (0) |
| **Pathological N stage** |  |  |  |
| pN0 | ypN0 | 33 (18) | 64 (36) |
| pN1 | ypN1 | 49 (27) | 18 (10) |
| pN2 | ypN2 | 7 (4) | 4 (2) |
| pN3 | ypN3 | 4 (2) | 0 (0) |
| **Molecular Subtype** |  |  |  |
| ER positive |  | 25 (14) |  |
| HER2 positive |  | 67 (37) |  |
| Triple negative |  | 87 (49) |  |
| **Chemotherapy Status** |  |  |  |
| Post Neoadjuvant Chemotherapy |  | 87 (49) |  |
| pCR |  | 50 |  |
| RCB1 |  | 5 |  |
| RCB2 |  | 13 |  |
| RCB3 |  | 8 |  |
| Not recorded |  | 11 |  |
| Treatment Naive |  | 92 (51) |  |
| **Histological Grade** |  |  |  |
| 1 |  | 5 (3) |  |
| 2 |  | 55 (31) |  |
| 3 |  | 119 (66) |  |
| **Lymph Node Procedure** |  |  |  |
| Sentinel Lymph Node Biopsy |  | 130 (73) |  |
| Up Front Axillary Clearance |  | 49 (27) |  |
| Completion Axillary Clearance |  | 22 |  |
| **Number of Lymph Nodes Analysed** |  |  |  |
| Reactive Nodes from Benign Cases |  | 23 |  |
| Uninvolved Nodes |  | 227 |  |
| Involved nodes |  | 104 |  |
| Macrometastasis |  | 71 |  |
| Micrometastasis |  | 26 |  |
| Isolated Tumor Cells (Post Neoadjuvant Chemotherapy) |  | 7 |  |

PDGFRβ mirrored reticulin staining, proving to be the most robust pan-FRC marker. It showed consistently strong staining in T cell zones, weak follicular dendritic cell (FDC) staining in germinal centers, and strong capsular fibroblast staining (**Figure 1**). SMA strongly marked FRC in the paracortex and capsule, with weak staining in parafollicular zones; it was absent in germinal centers. PDPN, strongly stained lymphatic endothelial cells and moderately labelled germinal center FDC, but did not highlight T zone FRC. Integrin β1 was restricted to mature blood vessels and the lymph node capsule. FAP staining proved technically challenging despite extensive optimization. The high temperatures required to retrieve antigen and achieve a signal damaged the tissue, and therefore FAP was not pursued further.

**Figure 1.**
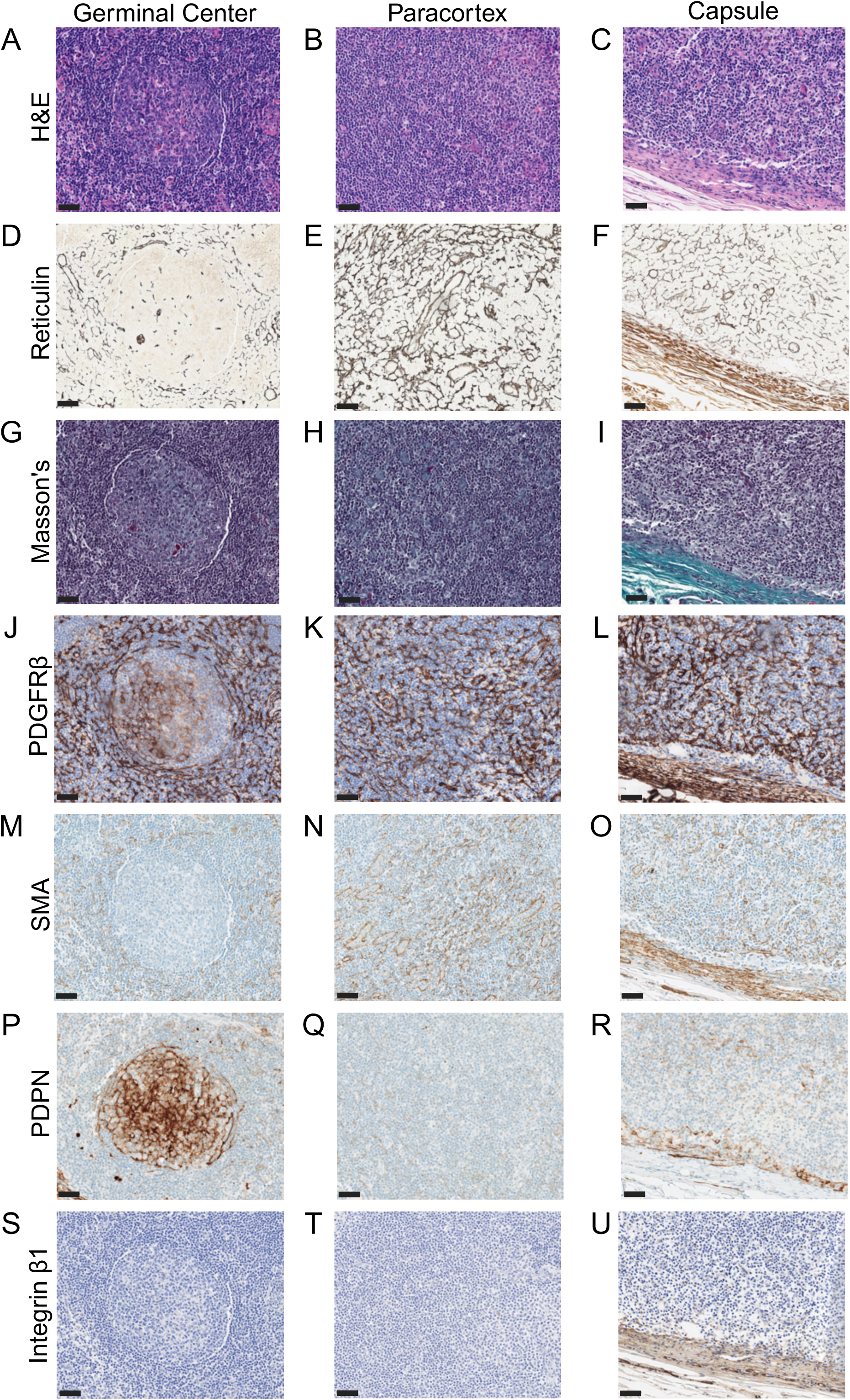
Optimized IHC Markers of Human FRC Subsets. These representative photomicrographs show optimized IHC staining of reactive human lymph node tissue from patients with benign disease (n = 23 nodes; scale bars, 50 µm; all images at x20 magnification). (A-C) The reticular network was not visible by H&E staining in the germinal centers (A), paracortex (T cell zone; B) or capsule (C). (D-F) Reticulin staining highlighted the reticular collagen network weakly in germinal centers (D) but strongly in the paracortex (E). The capsule (F) demonstrated a distinct reticulin staining pattern, characterized by thicker, wavier fibers. (G-I) Masson’s trichrome did not stain the FRC network in germinal centers (G) or the T cell zone (H), but stained the mature, capsular collagen blue (I). (K-L) PDGFRβ stained FRC in the paracortex (K), with weak FDC staining in germinal centers (J) and strong fibroblast staining in the capsule (L). This pattern mirrored reticulin staining (D-F). (M-O) SMA was absent in germinal centers (M), but stained FRC in the paracortex (N) and the capsule (O). (P-R) PDPN stained FDC in germinal centers (P) and lymphatic endothelial cells (R), but did not label FRC in the T cell zone (Q). (T-U) Integrin β1 staining was restricted to mature blood vessels and the capsule (U), with no staining of FDC (S) or FRC (T).

### Quantitative Metrics Capture FRC Network Architecture in Human Lymph Nodes

Whole slide images were scanned and representative 250,000 µm² regions of interest (ROI) were chosen from PDGFRβ-stained T cell zones. ROI were chosen from reactive nodes from patients with benign disease, uninvolved nodes from patients with BC, residual lymphoid tissue from nodes with smaller metastatic tumor deposits (< 2 mm in maximal extent), areas of NACT-induced fibrosis and areas of metastatic tumor (**Figure 2**). Since subtle alterations in FRC topology could potentially have large implications for tissue function, we needed a precise, unbiased approach to quantify changes in response to BC and/or NACT. The Workflow Of Matrix BioLogy Informatics (TWOMBLI) is an image analysis pipeline designed to quantify a broad range of parameters in the extracellular matrix in normal and pathological tissue ^27^. Due to the intimate association between FRC and the reticulin network, we were able to repurpose this pipeline to analyze FRC network parameters ^7^. TWOMBLI analysis showed that the FRC network in reactive lymph nodes (benign controls; clinico-pathological characteristics shown in **Supplementary Table 1**) has low lacunarity (median = 6, interquartile range (IQR) = 5-7.8), moderate branching (median branchpoints normalized by fiber length = 0.066, IQR = 0.059-0.068), consistent FRC size (median width = 1.5 µm, IQR = 1.3-2.0); median fiber length = 24 µm, IQR 23-34) and are poorly aligned (median alignment = 0.053, IQR = 0.04-0.077). Interestingly, these network features were not significantly correlated with patient age (**Supplementary Table 2**); however, this finding should be interpreted with caution due to the small sample size and the use of reactive nodes, which may not accurately reflect the uninflamed aging process.

**Figure 2.**
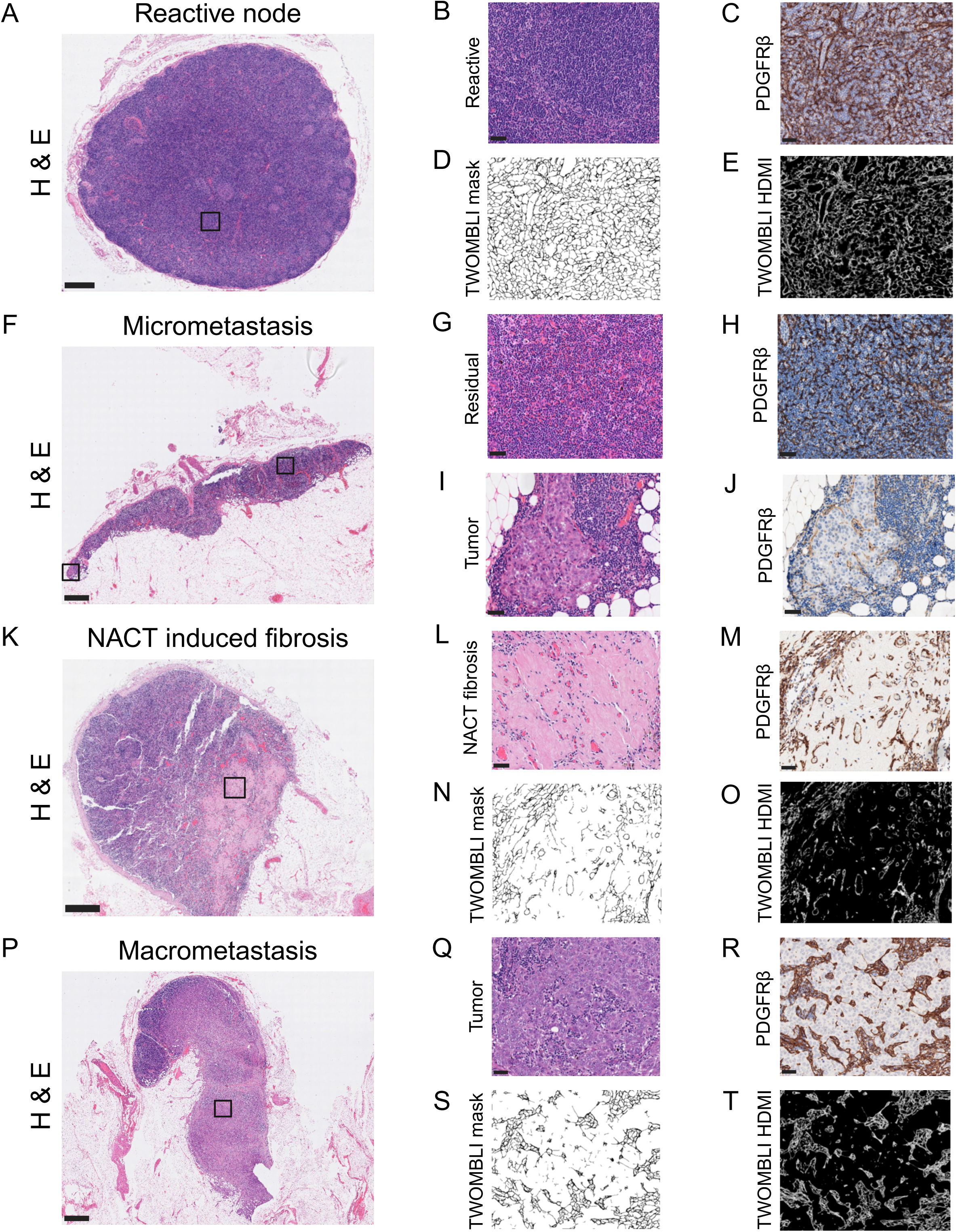
Quantitative Metrics Capture FRC Network Architecture in Human Lymph Nodes. ROI were selected for TWOMBLI analysis from reactive benign control (n = 23); uninvolved (n = 214); tumor-infiltrated (n = 75); and fibrotic (n = 16) nodes, as well as residual lymphoid tissue in metastatic nodes (n = 72). (A, F, K and P) The reticular network was not apparent on H&E-stained images (bird’s eye view; scale bars, 1mm). High-magnification (x20) images of boxed areas, each an example of analyzed ROI, are shown in B, G, I, L and Q (scale bars, 50 µm). (C, H, M and R) PDGFRβ highlighted a dense FRC network in reactive benign control and uninvolved nodes (C), as well as in residual lymphoid tissue in nodes containing small tumor deposits that were at least 500µm away (H; images at x20 magnification; scale bars, 50 µm). (J and R) In metastatic nodes, the small (J) and large (R) tumor deposits showed a disrupted, stretched FRC network (images at x20 magnification; scale bars, 50 µm). (M) In lymph nodes where tumor had been eradicated by neoadjuvant chemotherapy, leaving residual fibrosis, PDGFRβ highlighted only a few remaining, aligned FRC within this area (image at x20 magnification; scale bar, 50 µm). (D, N and S) From each PDGFRβ-stained ROI, TWOMBLI masks were generated to extract matrix features. This highlighted increased lacunarity in ALN containing large tumor deposits (S) compared to reactive nodes (D). (E, O and T) TWOMBLI masks were then processed and thresholded to calculate HDMI. This showed reduced HDMI in post-NACT fibrosis (O) compared to reactive nodes (E).

### Treatment-Naïve TNBC and NACT Remodel the FRC Network in Uninvolved ALN

Multivariate linear analysis of each TWOMBLI output from uninvolved nodes removed from BC patients against the clinico-pathological variables (**Table 1**) revealed statistically significant associations (**Supplementary Table 3**). Number of endpoints in the reticular network yielded the highest R score (0.127, p = 0.000007), with molecular subtype (p = 0.0006) and tumor burden in the axilla (p = 0.01; explained below) as significant predictors. Fiber width (p = 0.00006), number of branchpoints (p = 0.00009), lacunarity (p ≤ 0.0001), fiber length (p = 0.00007), high-density matrix intensity (HDMI; p = 0.001) and hyphal growth unit (HGU; p = 0.05) were all significantly predicted by NACT exposure.

To assess network parameters alongside clinical variables in a more integrative manner, we performed a Principal Component Analysis (PCA) on the combined dataset (**Figure 3A-C**). The first principal component (PC1) accounted for 26.09% of the total variance, predominantly driven by lacunarity (loading = -0.506), followed by branchpoints (loading = 0.478), and NACT (loading = 0.403) (**Supplementary Table 4** and **Supplementary Figure 1**). This suggests that PC1 captures variability driven by both network structural features and clinical context. Other PCs incorporated additional clinical and network parameters, indicating complex multi-variable relationships across topological and clinical parameters (**Supplementary Table 4**). Strikingly, the benign control lymph nodes always clustered separately from uninvolved ALN from BC patients, suggesting cancer-specific changes to ALN structure.

**Figure 3.**
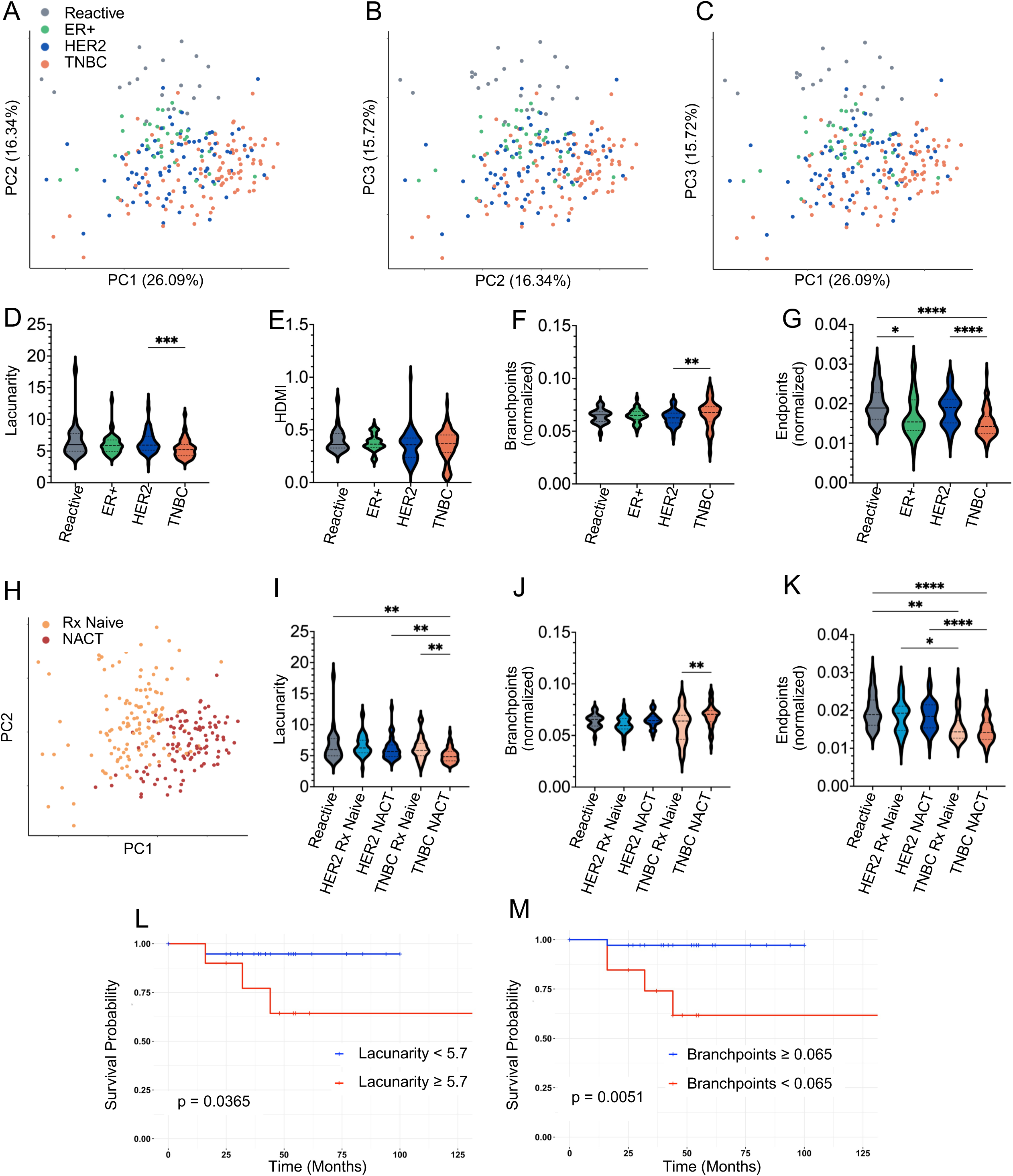
Treatment-Naïve TNBC and NACT Remodel the FRC Network in Uninvolved ALN. (A-C) PCA was performed to integrate the TWOMBLI derived FRC network features and clinico-pathological variables obtained from reactive, benign control nodes (n = 23) and uninvolved ALN (ER+: n = 40; HER2: n = 78; and TNBC: n = 96). Each data point represents the median of 8 ROI from each node. PC1 accounted for 26.09% of the total variance, predominantly driven by lacunarity (loading = -0.506), followed by branchpoints (loading = 0.478), and then NACT (loading = 0.403). PC2 and PC3 contributed 16.34% and 15.72% respectively to variance. See also Figure S1 and Table S2. Benign, reactive control nodes always clustered separately to uninvolved ALN from BC patients. (D-G) Violin plots showing molecular subtype-specific differences in FRC network lacunarity (D), HDMI (E), number of branchpoints (F) and number of endpoints (G) in uninvolved ALN *p ≤ 0.05, **p ≤ 0.01, ***p ≤ 0.001, ****p ≤ 0.0001 (Kruskal-Wallis test). Median with interquartile range, with minimum and maximum. See also Figure S2. (H) PCA showing effect of NACT on uninvolved nodes from all BC molecular subtypes (treatment naïve: n = 126 ALN; NACT: n = 111 ALN). Each data point represents the median of the results from 8 ROI from each node. (I, J and K) Violin plots illustrating the impact of NACT on FRC network lacunarity (I), number of branchpoints (J) and number of endpoints (K) in uninvolved ALN, stratified by BC molecular subtype (reactive: n = 23 nodes, HER2 treatment-naïve: n = 39 nodes, HER2 post-NACT: n = 39 nodes, TNBC treatment-naïve: n = 29 nodes, TNBC post-NACT: n = 67 nodes). Each data point represents the median of 8 ROI from each node. *p ≤ 0.05, **p ≤ 0.01, ***p ≤ 0.001, ****p ≤ 0.0001 (Kruskal-Wallis test). Median with interquartile range, with minimum and maximum. (L and M) Kaplan–Meier survival analysis of TNBC patients with ypN0 disease (i.e. no residual disease in the axilla post-NACT) shows that patients with uninvolved ALN that show a lacunarity ≥ 5.7 (p = 0.0365) or branchpoints <0.065 (p = 0.0051) have a significantly worse overall survival (n = 49 nodes; Gehan-Breslow-Wilcoxon method).

Further, we observed topological alterations linked to BC subtype. Stratification of TWOMBLI results from both treatment naïve and post-NACT patients by BC molecular subtype revealed that FRC networks in uninvolved nodes from patients with TNBC had significantly reduced lacunarity (p ≤ 0.001; **Figure 3D**), unchanged HDMI (**Figure 3E**), increased branchpoints (p = 0.0048; **Figure 3F**) and fewer endpoints (p ≤ 0.0001; **Figure 3G**) compared to HER2 and reactive nodes, independently of treatment. Visualization of the PCA distribution by NACT exposure also revealed a clear segregation between treatment-naïve and NACT-exposed patients (**Figure 3H**). Subsequent faceting of the PCA by both molecular subtype and NACT exposure further demonstrated distinct separation between uninvolved and reactive control nodes (**Supplementary Figure 2**). However, the separation was more pronounced in TNBC than HER2 positive disease and was even greater in nodes from patients who had received NACT, indicating that both tumor subtype and therapy exposure contribute to network remodeling. Further stratification of individual TWOMBLI parameters according to subtype and NACT exposure confirmed these patterns. FRC networks from NACT-exposed TNBC patients displayed significantly decreased lacunarity (p ≤ 0.01; **Figure 3I**), increased branchpoints (p ≤ 0.01; **Figure 3J**) and decreased endpoints (p ≤ 0.0001; **Figure 3K**; **Supplementary Figure 2)**.

Collectively, these findings indicate that in treatment-naive TNBC patients, the FRC network in uninvolved ALN is more compact and highly branched. Interestingly, exposure to NACT induces comparable alterations throughout the ALN basin. Notably, Kaplan–Meier analysis of patients with TNBC and no axillary nodal involvement following NACT (pathological nodal stage 0, ypN0) demonstrated that the presence of a denser FRC network, characterized by lower lacunarity (p = 0.0365, **Figure 3L**) and a higher number of branchpoints (p = 0.0051, **Figure 3M**), correlated with improved survival.

### Metastatic Tumor Deposits Drive Greater FRC Network Disruption Within Residual Nodes

In order to test the effect of BC on the FRC network within a single node as well as across the ALN chain, we repeated our analysis including ROI of areas of residual lymphoid tissue in involved nodes from patients of all molecular subtypes, both treatment naïve and post-NACT. Multivariate analysis revealed lacunarity as the parameter most significantly influenced by clinical factors (R score 0.157, p = 2×10⁻⁹) with the size of metastasis (i.e. ITCs; micrometastasis; or macrometastasis), NACT exposure and molecular subtype as the most significant predictors (**Supplementary Table 5**). Direct comparison of TWOMBLI outputs from treatment-naive and post-NACT nodes confirmed the presence of a denser, more branched network demonstrated in the uninvolved nodes following NACT (**Supplementary Figure 3**). Furthermore, stratification by molecular subtype also confirmed a significant effect of TNBC on topology.

PCA again demonstrated that across all subtypes, reactive control nodes from benign patients formed distinct clusters compared to BC patient-derived ALN (**Figure 4A-I**). Moreover, within each molecular subtype, metastatic ALN containing residual lymphoid tissue clustered separately to uninvolved nodes. This indicates that the effects of BC on the stromal network are more pronounced in the residual lymphoid tissue that sits adjacent to tumor deposits in metastatic ALN. Interestingly, lacunarity had the highest loading on PC1 (0.410), which accounted for 26.62% of the variance in the data (**Supplementary Table 6** and **Supplementary Figure 1**). This aligns with the results of the multivariate analysis (**Supplementary Table 5)**. Furthermore, PC1, PC2, and PC3 all contributed to meaningful separation of the data (**Figure 4J-L**). Taken together, these results confirm that the presence of TNBC and exposure to NACT induce a more compact FRC network in the residual lymphoid tissue of involved nodes.

**Figure 4.**
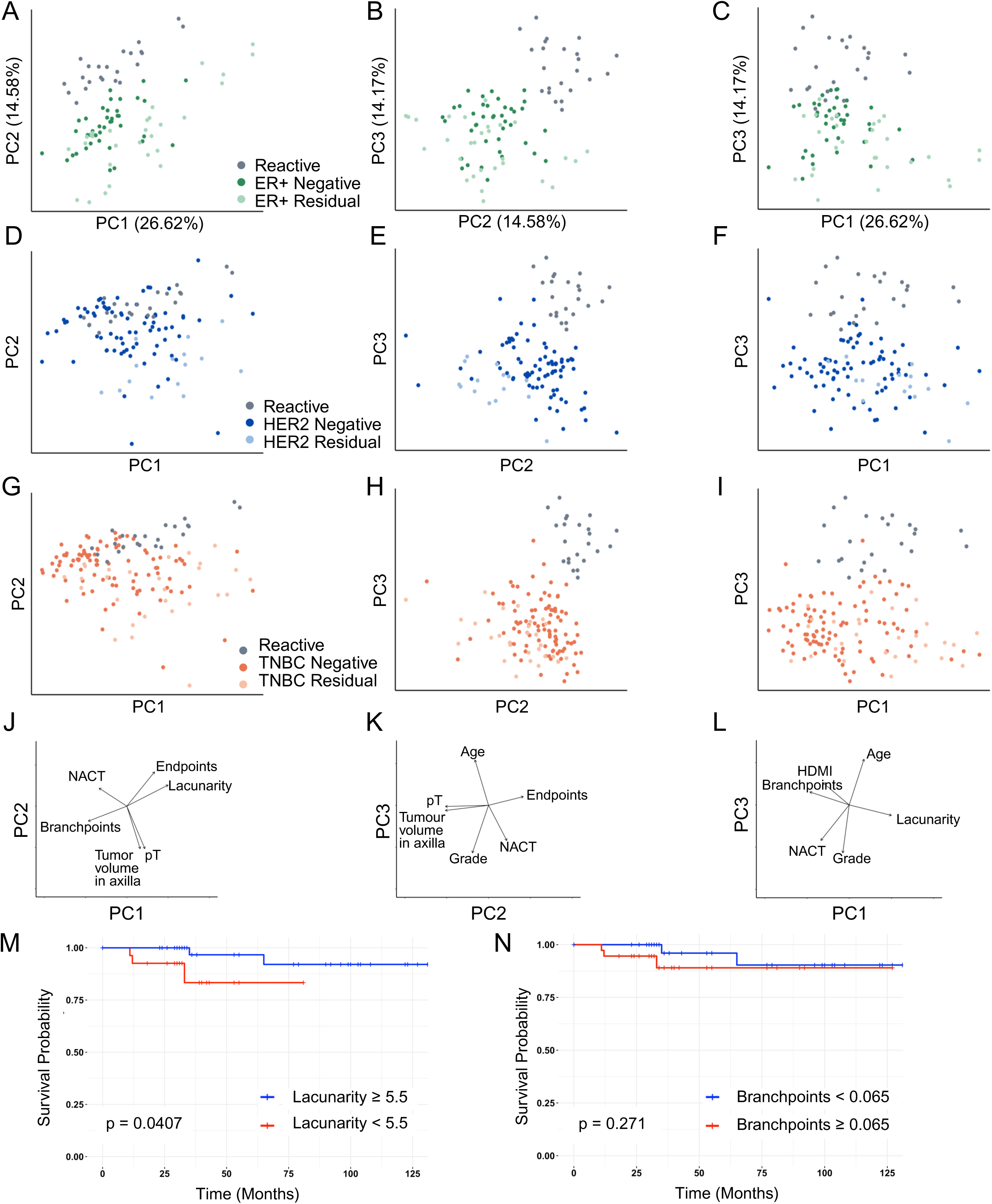
Metastatic Tumor Deposits Drive Greater FRC Network Disruption Within Residual Nodes. PCA, stratified by BC molecular subtype, was performed to integrate TWOMBLI derived FRC network features and clinico-pathological variables from reactive benign nodes (n = 23), uninvolved ALN and residual lymph node tissue from involved ALN. PC1, PC2, and PC3 contribute 26.62%, 14.85%, and 11.47% of the variance, respectively. Each data point represents the median of 8 ROI per node. See also Figure S1, Figure S3 and Table S4. Once again, reactive benign control nodes clustered distinctly from BC patient-derived nodes. (A, B and C) ALN from patients with ER positive BC (ER+ uninvolved: n = 40 nodes, ER+ residual: n = 29 nodes). (D, E and F) ALN from patients with HER2 positive disease (HER2 uninvolved: n = 78 nodes, HER2 residual: n = 15 nodes). (G, H and I) ALN from patients with TNBC (TNBC uninvolved: n = 96 nodes, TNBC residual: n = 28 nodes). (J, K and L) Loading plots illustrating the contribution of the top six individual features to the first three principal components. (M and N) Kaplan–Meier survival analysis of patients with pN1 or ypN1 disease, of any molecular subtype, showed that patients with uninvolved ALN with a lacunarity <5.5 have a significantly worse overall survival (p = 0.0407) but that branchpoints ≥ 0.065 (p = 0.271) is not a significant predictor (n = 82 nodes; Gehan-Breslow-Wilcoxon method).

We specifically investigated the impact of tumor metastasis in ALN on the FRC topology of uninvolved nodes within the same chain and linked these data with patient survival. In patients with metastatic BC involving one to three lymph nodes (pathological nodal stage 1, pN1 (treatment-naïve) and/or ypN1 (post-NACT)), a more compact FRC network, indicated by reduced lacunarity, in the uninvolved nodes was significantly associated with poorer overall survival, independent of treatment status (p = 0.0407; **Figure 4M**). However, in this group of patients, number of branchpoints was not correlated with survival (p = 0.271; Figure 4N).

### Chemotherapy-Induced Fibrosis Replaces Reticular Network Architecture

In reactive, benign control lymph nodes the delicate reticulin/FRC network spans the entire node (**Figures 5A-D**). In contrast, NACT produces wedge-shaped areas of fibrosis in which tumor cells are replaced by dense eosinophilic collagen and angiogenic vessels **(Figure 5E)**. This fibrosis clinically indicates a response to NACT ^28^. However, the nature of this fibrosis and its effect on the FRC network have not been previously characterized. Reticulin staining of these areas revealed abundant mature collagen composed of thicker, wavy, argyrophilic fibers that replaced the normal reticular network (**Figure 5F**). These fibers were similar to those found within the lymph node capsule and also showed blue staining with Masson’s trichrome (**Figure 5G**), indicating the presence of mature collagen. PDGFRβ immunostaining demonstrated complete destruction of the network with fewer, scattered remaining FRC (**Figure 5H**). TWOMBLI analysis revealed increased FRC alignment (p = 0.0210; **Figure 5M**), non-significant changes in HDMI (**Figure 5N**) and a significant reduction in FRC length (p = 0.002; **Figure 5O**) in these fibrotic areas. However, TWOMBLI is not optimized for assessing alignment, particularly in areas where the network has been destroyed. Therefore, the same ROIs were additionally analyzed using CurveAlign ^29^, which showed a trend towards increased FRC alignment (p = 0.09; **Figure 5P**) in areas of post-NACT fibrosis when compared to matched, uninvolved ALN, but this did not reach statistical significance.

**Figure 5.**
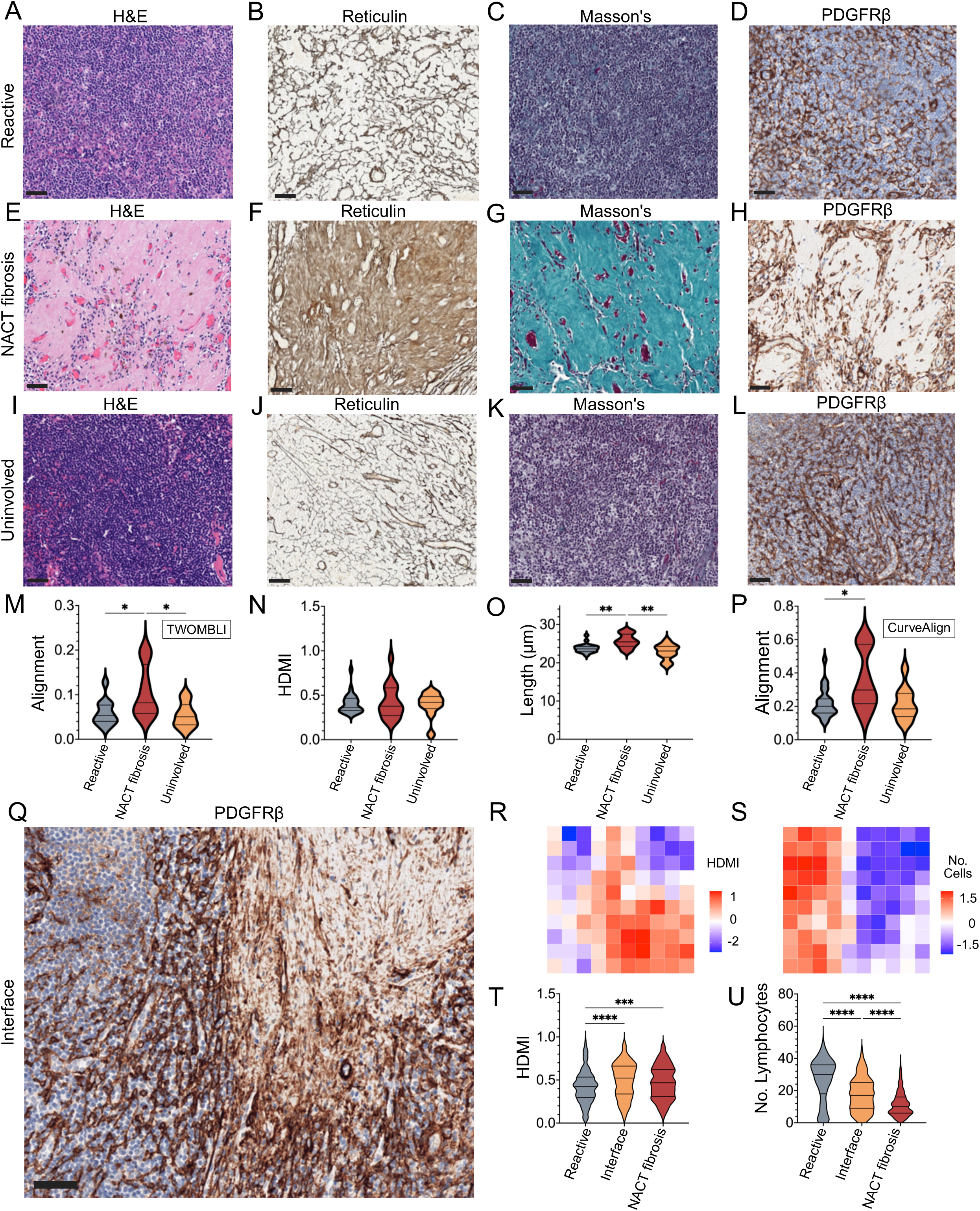
Chemotherapy-Induced Fibrosis Replaces Reticular Network Architecture. (A-D) Representative photomicrographs showing ROI from reactive, benign control lymph nodes (n = 23; x20 magnification, scale bars, 50 µm). The reticular network was not visible by H&E staining (A), but was highlighted by reticulin staining (B). Masson’s Trichrome (C) showed no mature collagen in the reactive lymph node parenchyma. PDGFRβ stained the delicate FRC network (D). (E-H) Representative photomicrographs showing ROI from areas of chemotherapy-induced fibrosis (n = 16 nodes; x20 magnification, scale bars, 50 µm). H&E staining (E) showed dense eosinophilic collagen with sparse lymphocytes. Reticulin staining (F) showed complete replacement of the normal reticular network by thicker argyrophilic fibers. Masson’s trichrome staining (G) demonstrated abundant mature (blue) collagen. PDGFRβ immunohistochemistry (H) revealed complete destruction of the FRC network with few remaining FRC. (I-L) Representative photomicrographs showing ROI from areas of uninvolved ALN that were removed from the same patients as those containing NACT-induced fibrosis (n = 16 nodes; x20 magnification, scale bars, 50 µm). H&E staining (I) showed only lymphoid tissue, with no areas of fibrosis and no evidence of tumor deposits. Reticulin staining (J) showed a normal reticular network. Masson’s trichrome stain (K) showed no mature collagen. PDGFRβ immunostaining (L) revealed a preserved FRC network. (M-P) Violin plots comparing FRC alignment (M and P), HDMI (N) and length (µm) across reactive lymph nodes (n = 23) to areas of chemotherapy-induced fibrosis and matched uninvolved (negative) ALN from the same patient (n = 16). FRC alignment calculated using both TWOMBLI software (M) and CurveAlign software (P;see methods for details). Each data point represents a median of the results from 4 ROI from each node. *p ≤ 0.05 (Kruskal-Wallis test). Median with interquartile range, with minimum and maximum. (Q) Representative photomicrograph of a PDGFRβ-stained ROI showing the interface between chemotherapy-induced fibrosis and the residual FRC network (n = 13 nodes; x20 magnification, scale bars, 50 µm). (R and S) Examples of heatmaps of TWOMBLI analysis generated from ROI in (Q) illustrating the gradual increase in HDMI (R) and decrease in lymphocyte cell number (S) across the transition from an area of residual lymph node to an area of NACT-induced fibrosis. (T and U) Violin plots of the aggregated heatmap data comparing HDMI (T) and number of lymphocytes (U) across areas of residual lymph node, interface and areas of NACT-induced fibrosis (n = 13 nodes). ***p ≤ 0.001, ****p <0.0001 (Kruskal-Wallis test). Median with interquartile range, with minimum and maximum.

PDGFRβ staining at the interface between chemotherapy-induced fibrosis and the remaining FRC network showed the gradual transition from destroyed to disrupted to intact FRC network architecture (**Figure 5Q**). For high-resolution quantification, each ROI at the interface was subdivided into 100 tiles, which were individually analyzed using TWOMBLI. These tiles were then reconstructed into heatmaps to visually represent the gradual changes in network parameters across the residual node, interface and fibrotic area (examples shown in **Figures 5R and 5S**). Aggregated heatmap data showed the highest average HDMI values in the interface region (p ≤ 0.0001; **Figure 5T**), and a gradual decrease in cell (lymphocyte) number from the residual node through the interface to the fibrotic zone (p ≤ 0.0001; **Figure 5U**).

### FRC Network Topology Correlates with Tumor Burden and Clinical Outcome in Metastatic ALN

As anticipated, survival analysis of this cohort demonstrated a poorer prognosis for patients with a higher nodal stage (**Figure 6A**), as well as those with TNBC and a higher tumor stage (**Supplementary Figure 4**). However, traditional nodal stage incorporates only the number of positive nodes and not the size of deposits, which we have shown to be highly relevant to network topology. Furthermore, in the clinical setting, axillary tumor burden is routinely quantified only in post-NACT samples and not in treatment naïve disease ^30^. In order to compare these two patient cohorts, we adapted the formula used to calculate the residual cancer burden score to estimate axillary tumor burden (see **Methods**) ^30^. Stratification using this metric revealed that patients with a tumor volume ≥ 20 had significantly worse outcomes (p = 0.0075; **Figure 6B**). As tumor cells invade ALN, they progressively distort the FRC network, quantified by a significant increase in lacunarity correlated with a decrease in HDMI (**Figure 6C-G).** We observed that the degree of this network distortion is correlated to the size of the metastatic deposit, rather than molecular subtype or NACT exposure (**Figure 6C-G**). Notably, large nests of tumor resulting in high network lacunarity (≥ 12) was associated with significantly poorer survival (p = 0.043, **Figure 6H**). More surprisingly, in patients with BC of all molecular subtypes and treatment status, increased FRC alignment in metastatic deposits improved survival (p = 0.0205, **Figure 6I-K**).

**Figure 6.**
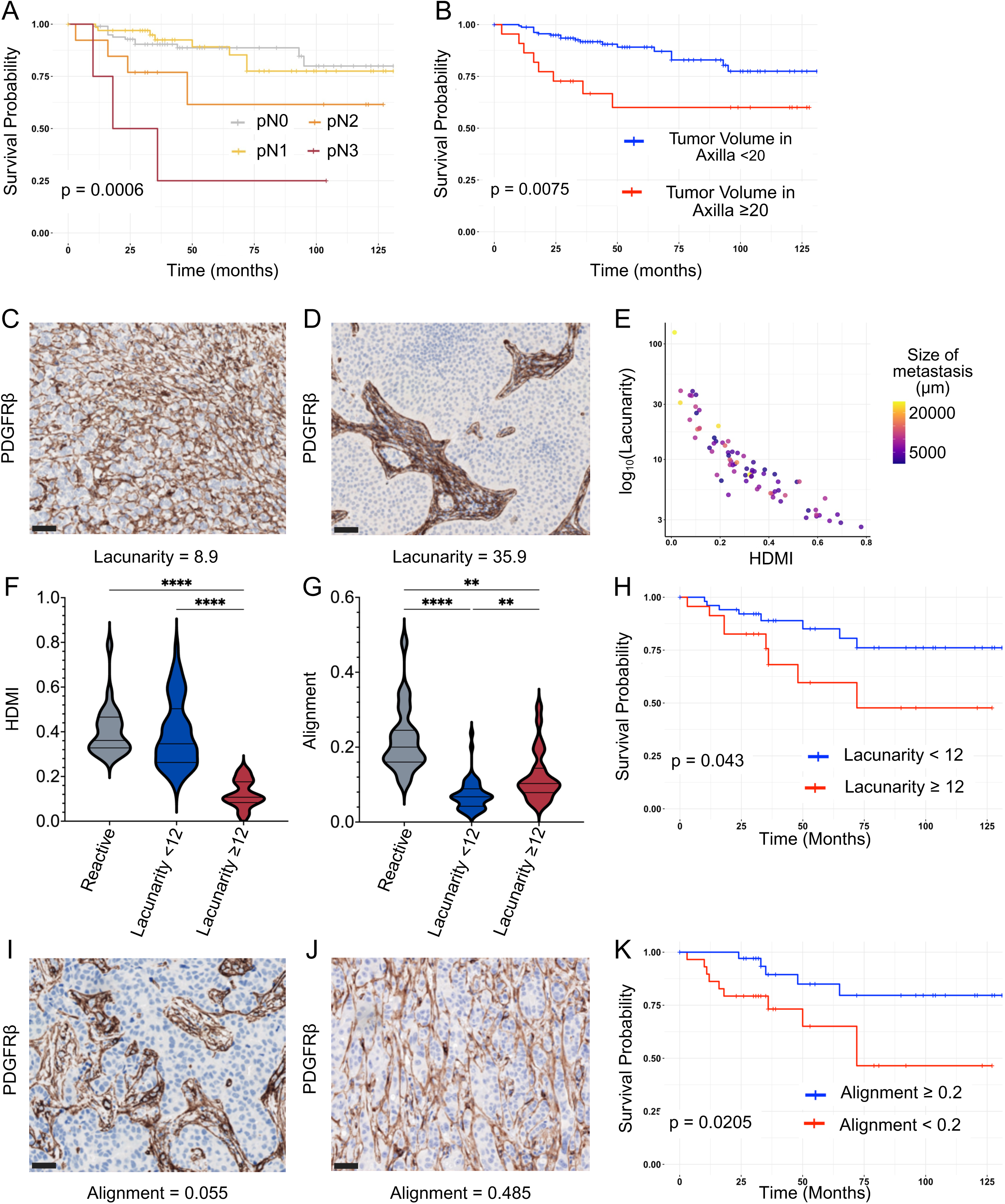
FRC Network Topology Correlates with Tumor Burden and Clinical Outcome in Metastatic ALN. (A and B) Kaplan–Meier survival curves showed reduced survival in patients with higher nodal stage (A; p = 0.0006) and axillary tumor volume ≥ 20 (B;p = 0.0075; n = 179 patients; Gehan-Breslow-Wilcoxon method). See also Figure S4. (C and D) Representative photomicrographs showing ROI of macrometastases with different lacunarities in ALN stained with PDGFRβ (x20 magnification, scale bar, 50 µm). (E) Scatterplot showing the inverse correlation between TWOMBLI derived lacunarity and HDMI in metastatic ALN. Data point color reflects metastatic tumor diameter (µm); larger metastases are associated with higher lacunarity and reduced FRC network density (n = 75 nodes; values for lacunarity and HDMI represent a median of the results from 8 ROI in each node). (F and G) Violin plots comparing FRC network HDMI (F) and alignment (G) in involved ALN, stratified according to lacunarity (involved ALN with lacunarity <12: n = 52; involved ALN with lacunarity ≥12: n = 23) to reactive benign control nodes (n = 23). Each data point represents a median of the results from 8 ROI from each node. **p ≤ 0.01, ****p ≤ 0.0001 (Kruskal-Wallis test). Median with interquartile range, with minimum and maximum. (H) Kaplan–Meier survival analysis, stratified by lacunarity in metastatic nodes, showed that patients with positive nodes showing high lacunarity have significantly poorer survival compared to those with low lacunarity (p = 0.043; n = 64 patients; Gehan-Breslow-Wilcoxon method). (I and J) Representative photomicrographs showing ROI of a macrometastasis with an alignment of 0.055 (I, calculated using CurveAlign software), as opposed to one with an alignment of 0.4 (J), in ALN stained with PDGFRβ (x20 magnification, scale bar, 50 µm). (K) Kaplan–Meier survival analysis, stratified by alignment in metastatic nodes, showed that patients with involved nodes showing an FRC alignment <0.2 (calculated using CurveAlign software) have significantly poorer survival compared to those with higher FRC alignment (p = 0.0205; n = 64 patients; Gehan-Breslow-Wilcoxon method).

## DISCUSSION

By optimizing robust IHC markers for human FRC, we have shown how different subsets of FRC localize within the ALN of BC patients. We confirmed PDGFRβ as a pan-FRC marker that highlights FRC in all of the microanatomical ALN compartments, where different immune populations reside and traffic in both health and disease ^22^. Interestingly, our data show that, in contrast to murine lymph nodes, where it stains FRC in the T cell zones, human PDPN is highly expressed predominantly by germinal center FDC and lymphatic endothelial cells ^14^. In addition, SMA expression is more prominent in human tissue, where it is seen primarily in the paracortex, than in mice, where it typically stains weakly in germinal centers, increasing only upon activation. It is possible that this difference in staining patterns, rather than being an inherent variation between species, is age-related, since most mouse models use young, immunologically naïve mice, whereas in humans, even ‘normal’ nodes will have undergone repeated cycles of antigen exposure, immune activation and resolution before being removed and examined. Moreover, PDPN expression has been observed in FRC within tertiary lymphoid structures in human non-small cell lung cancer, indicating that its expression may be context-dependent ^31^. Regardless, this biological variablity could compromise on-going efforts at clinical translational, and as such, should be investigated further.2

Because of the intimate association between FRC and collagen in lymph nodes ^32^, we were able to repurpose TWOMBLI as an unbiased and quantitative tool to map the human FRC network in different pathophysiological conditions ^27^. By integrating TWOMBLI derived parameters with clinical data through multivariate analysis, we were able to clearly visualize FRC network changes in a translationally relevant context. Although we focused on BC, our approach is applicable to other malignancies and pathologies.

We have demonstrated that the FRC network becomes denser and more highly branched in treatment naïve TNBC, an aggressive, hard-to-treat BC subtype. We postulate that the denser FRC network seen in these patients may restrict the flow of lymph through the reticular network in the ALN parenchyma, shunting fluid through the SCS and into the efferent vessels. This change in fluid flow could likely have two clinically relevant effects. Firstly, the pooling of fluid in the SCS may provide an opportunity for cancer cells to extravasate and accumulate, leading to the development of metastatic tumor deposits ^33^. Secondly, controlled flow through the lymph node parenchyma is critical for antigen sampling by tissue resident antigen-presenting cells. Without these interactions the ability of the node to generate effective immune responses against tumor antigens and pathogens would be compromised ^10^. Our data show these changes trend towards a poorer prognosis in patients with treatment naïve TNBC, but this result did not reach statistical significance. This is most likely due to the fact that it is hard to collect tissue from treatment naïve patients, since most TNBC patients receive NACT at present ^34^.

Interestingly, these topological changes were evident in both patients without nodal metastasis and in uninvolved ALN from patients with metastatic deposits elsewhere in the axillary chain. Previous mouse studies have shown that factors secreted by primary melanoma travel to the draining lymph nodes, promoting FRC proliferation and a shift toward a cancer-associated fibroblast (CAF)-like phenotype ^35^. Thus, it stands to reason that soluble factors originating in the primary breast tumor could be responsible for the changes seen in patients with pN0 disease. On the other hand, the presence of similar topological changes in uninvolved nodes from patients with nodal metastasis in other ALN (pN1/ ypN1) could be due to inter-nodal communication within the axillary chain through soluble factors. This makes it difficult to unpick precisely what is occuring in patients who received NACT, since these patients could have had metastatic disease in their axillary nodes which were eradicated before surgery. In other words, inter-nodal communication cannot be entirely excluded in ypN0 cases. This confounding effect is not seen once metastases are established in the axillary chain, however. In the uninvolved nodes from patients with pN1 or ypN1 disease of all subtypes, a reduction in lacunarity was associated with poorer prognosis, regardless of treatment status. This is clinically significant as treatment decisions regarding the extent of axillary surgery in the node-positive cohort are often challenging ^3^. Therefore, any biomarker that can predict whether SLN have switched from ‘helping’ (immunocompetent) to ‘harming’ (immune-exhausted) a patient would be invaluable.

Biologically, TNBC is thought to be more immunogenic than other BC subtypes. A subset of TNBC tumors are lymphocyte-predominant (i.e. contain tumor infiltrating lymphocytes amounting to at least 30% of the tumor) and ALN of patients with TNBC more often contain a greater number of GC than other BC subtypes ^36,37^. Despite this, CPI are only effective if NACT is given concurrently to augment the anti-tumor immune response ^38^. Nevertheless, only 50% of TNBC patients achieve a complete pathological response after NACT, and only a further 10% benefit from the addition of a CPI to neoadjuvant treatment regimens ^39^. We have shown that the FRC network of uninvolved ALN is significantly denser and more highly branched post-NACT, and that this was more pronounced in TNBC patients. Intriguingly, in contrast to the treatment naïve patients, this change in FRC topology was associated with an improved survival in TNBC patients with no residual metastasis in the axilla. This suggests that the same alterations in fluid flow, with shunting into the SCS, may be beneficial in this cohort, potentially by facilitating drug-induced killing of small subcapsular tumor deposits.

In metastatic ALN where chemotherapy eradicates large tumor deposits, it concurrently obliterates the FRC network, replacing it with dense, mature fibrotic collagen structures containing a few, highly aligned FRC and very occasional lymphocytes. These structural changes could have important functional consequences for impeding immune cell trafficking and activation, and this in turn might help to resolve whether patients who have a complete pathological response after NACT should have a targeted, rather than a complete, axillary dissection ^40^. Moreover, since the addition of CPI to NACT results in mixed responses, understanding all of the factors that could influence treatment response is crucial. We aim to confirm and further characterize this in a larger cohort of patients in the future.

Simulations indicate that the FRC network remains functionally resilient despite the loss of up to 50% of its structure ^41^. We have shown that even though the FRC network is maintained as BC invades into a node, its topology is highly distorted. Furthermore, remodeling of the FRC network to a denser, more branched topology was observed in residual lymphoid tissue that was situated > 500 µm from metastatic deposits within a node. These changes were similar to those seen in the uninvolved nodes but to a greater extent. Simple diffusion of the tumor-derived soluble factors through a whole lymph node is unlikely to be responsible for these changes as the distances are too great ^42,43^. It seems more plausible that the tumor-derived soluble factors driving these intranodal changes are transported through the altered conduit network.

As axillary tumor burden increased in all of the BC patients in our cohort, survival significantly decreased. This reinforces the prognostic value of both the number of nodal deposits and the size of each deposit, in both the treatment naïve and post-NACT context. Furthermore, we have shown that the extent of FRC network distortion within the metastatic deposit correlates with the size of the deposit rather than the BC subtype or other clinical features, suggesting that physical disruption is the primary driver. Once a tumor deposit exceeds a lacunarity of 12, survival worsens significantly. This size threshold could indicate a critical point at which the altered flow of fluid through the FRC network compromises the ability of the node to mount effective responses, thereby adversely affecting survival. Conversely, increased FRC alignment in tumor deposits is associated with a better prognosis, which may reflect more controlled flow through the node parenchyma. These hypotheses warrant further investigation using *in vivo* and *in silico* models to fully elucidate the mechanisms by which FRC network architecture influences fluid flow and immune function. We intend to use the REPLICANT model, a unique *ex vivo* perfusion system for human lymph nodes, to study fluid flow in real time and identify key molecular drivers and/or clinically relevant biomarkers ^44–46^.

We have shown, that changes in FRC topology can be quantified using unbiased computational approaches in human ALN using IHC, and that these changes predict prognosis in BC. By integrating this with real time flow dynamics, we are now positioned to mechanistically disentangle these complex interactions in patient-derived samples in translationally relevant context in future.

### Limitations of the study

None of the patients in this cohort received CPI, as these therapies were only recently introduced into routine clinical practice and therefore, outcome data for these samples would amount to less than 5 years^6^. Given the growing use of immunotherapy, particularly in TNBC, future studies should explore how combination therapies involving CPI and NACT impact FRC network architecture and function.

In addition, although this was a longitudinal study with ALN tissue samples collected from patients at various stages of disease and with follow-up data extending up to 10 years, the availability of sequential samples from individual patients was limited. Despite this, we were able to identify meaningful changes in the FRC network over time. Future prospective studies with longitudinal sampling after surgical resection of the primary tumor would enhance our understanding of the permanence or potential reversibility of these stromal changes.

## RESOURCE AVAILABILITY

### Lead contact

Further information and requests for resources should be directed to and will be fulfilled by the lead contact, Kalnisha Naidoo.

### Materials availability

This study did not generate new unique reagents.

### Data and code availability

All data reported in this paper will be shared by the lead contact upon request. This paper does not report original code.

Any additional information required to reanalyze the data reported in this paper is available from the lead contact upon request.

## ACKNOWLEDGMENTS

We would like to thank the patients for consenting to the use of their tissue for this project. This work was funded by Breast Cancer Now (Dame Vera Lynn Breast Cancer Now Clinical Research Training Fellowship 2023.06CRTF1643, A.L) and Cancer Research UK (RCCSCF-May22\100001, S.E.A).

## AUTHOR CONTRIBUTIONS

Conceptualization, K.N., S.E.A. and A.L.; methodology, A.L., S.D.C. R.L, J.G., V.L. and D.S.; investigation, A.L.; writing—original draft, A.L.; writing—review & editing, K.N. and S.E.A.; funding acquisition, A.L., K.N. and S.E.A.; resources, K.N., S.E.A., A.L., S.D.C and R.L.; supervision, K.N. and S.E.A.

## SUPPLEMENTAL INFORMATION

Document S1. Figures S1 – S4 and Tables S1 – S6.

## STAR*METHODS

### KEY RESOURCES TABLE

| REAGENT or RESOURCE | SOURCE | IDENTIFIER |
| --- | --- | --- |
| Antibodies |  |  |
| Platelet-derived growth factor receptor beta (PDGFRβ), RM303 | Invitrogen | Product code: MA5-33050 |
| Podoplanin, D240 | Cell Marque | Product code: 322M-10 |
| Smooth muscle actin (SMA), 1A4 | Cell Marque | Product code: 202M-90 |
| Integrin beta 1 (CD29), EP1041Y | Abcam | Product code: ab52971 |
| Fibroblast activation protein (FAP), EPR20021 | Abcam | Product code: ab207178 |
| Biological samples |  |  |
| FFPE human axillary lymph node samples | Patients, under appropriate ethical consent | N/A |
| Chemicals, peptides, and recombinant proteins |  |  |
| Xylene | ThermoFisher Scientific | Cat #X/0100/PB17 |
| 1% Potassium Permanganate | Sigma Aldrich | Cat#223468 |
| Sulphuric Acid | Sigma Aldrich | Cat#339741 |
| Oxalic Acid | Sigma Aldrich | Cat#194131 |
| Ferric Ammonium Sulphate | Sigma Aldrich | Cat#221260 |
| Formaldehyde | Sigma Aldrich | Cat#F8775 |
| Sodium Thiosulphate | Sigma Aldrich | Cat#217263 |
| Silver Nitrate | Sigma Aldrich | Cat#209139 |
| Ammonia | Sigma Aldrich | Cat#499145 |
| Ponceau De Xylidine (CI16150) | ThermoFisher Scientific | Product code: 10012450 |
| Glacial Acetic Acid | Sigma Aldrich | Cat#A6283 |
| Acid Fuchsin (CI42685) | Avantor Scientific | Cat#ICNA0215819725 |
| Light Green SF (CI42095) | Avantor Scientific | Cat#ICNA0215264125 |
| Molybdophosphoric Acid | Sigma Aldrich | Cat#221856 |
| Software and algorithms |  |  |
| Qpath (0.5.1-x64) | Bankhead et al. <sup>47</sup> | https://qupath.github.io/ |
| Image J Fiji | Schneider et al. <sup>48</sup> | https://imagej.net/software/fiji/ |
| TWOMBLI | Wershof et al. <sup>27</sup> | https://github.com/wershofe/TWOMBLI |
| Mathworks MATLAB | N/A | https://uk.mathworks.com/products/matlab.html |
| CurveAlign 5.0 | Liu et al. <sup>29</sup> | https://github.com/uw-loci/curvelets/releases/tag/5.0 |
| R (v4.3.1) | The R Project | https://www.r-project.org/ |
| GraphPad Prism (version 10.4.1) | N/A | https://www.graphpad.com/features |
| Other |  |  |
| Epredia™ SuperFrost Plus™ Adhesion slides | ThermoFisher Scientific | Product code: 10149870 |
| BOND-III Fully Automated IHC and ISH Staining System | Leica Biosystems | Product code: 21.2201 |
| Ventana BenchMark ULTRA IHC/ISH System | Roche | N/A |
| Glissandro 20SL Desktop Scanner | Objective Imaging | N/A |

## EXPERIMENTAL MODEL AND STUDY PARTICIPANT DETAILS

### Patient Cohort

Archival, FFPE human tissue samples were obtained through the Breast Cancer Immune, Drug and Gene (BRIDGE) Study (Research Ethics Committee No: 24/NW/0079). KCH pathology databases were searched to identify patients who had undergone a breast surgical excision, with either a SLNB, ALNC or both, for invasive BC between January 2014 and January 2024. A total of 179 patients were selected to ensure a representative cohort encompassing the three BC molecular subtypes (i.e. ER positive, HER2 positive and TNBC), as well as varying axillary (nodal) involvement and responses to NACT.

For each case, the following clinico-pathological data were collected from the electronic patient record: age, molecular subtype, pathological TNM stage, histological grade, NACT received, response to NACT, lymph node procedure performed, size of primary tumor, size of largest ALN metastasis and overall survival (**Table 1**). In order to calculate tumor burden in the axilla, we adapted the formula used to calculate residual cancer burden proposed by Symmans et al. ^30^: *tumor volume in the axilla = (1 − 0.75^LN^) d_met_)*, where LN is the number of positive lymph nodes and d_met_ is the diameter of the largest nodal metastasis. Overall survival was defined as time from diagnosis to death of any cause.

An additional 23 patients without a diagnosis of breast or any other cancer were also identified. These patients had undergone a lymph node biopsy or excision for non-malignant indications and were ultimately diagnosed with benign conditions, including follicular hyperplasia or reactive lymphadenopathy. These lymph nodes served as benign, reactive controls.

Diagnostic slides were reviewed by Histopathologists (AL and KN) to confirm ALN disease status and select appropriate blocks for immunohistochemical and digital analysis. In each case, where available, a lymph node containing a metastastic deposit and a negative node were chosen. In cases with both micrometastasis and macrometastasis, examples of both were selected. In patients who had SLNB followed by ALNC, nodes from both procedures were included. Negative nodes from patients who underwent ALNC were not ordered or labeled by nodal level during surgery, and therefore were considered representative of the entire axillary basin. Each tissue sample, together with the associated data, was de-identified before being released to researchers to ensure blinded downstream analysis.

## METHOD DETAILS

### Tinctorial Staining

Sections were cut at 3 µm thickness, then deparaffinised in xylene and rehydrated through graded alcohols and distilled water.

Reticulin fibers were visualised using a modified silver impregnation method. Slides were first oxidized in acidified potassium permanganate (equal parts 1% potassium permanganate and 0.3% sulfuric acid) for 7 min. After rinsing, sections were bleached in 1.5% oxalic acid until colorless (1 min), followed by immersion in 2.5% iron alum (ferric ammonium sulfate, 2.5g/100 mL distilled water) for 10 min. Fresh silver solution was prepared by combining 7mL 10% silver nitrate with 3 mL industrial methylated spirits (IMS), followed by dropwise addition of 0.88 ammonia until the initial precipitate dissolved, then adding 5 additional drops of ammonia. The silver solution was applied to each section for 1 min. Slides were then developed for 1 min using a formaldehyde-based reticulin developer and stabilized in 5% sodium thiosulfate (2 min). Between each staining step, slides were rinsed with distilled water.

Masson’s trichrome staining was performed using a celestine blue–haematoxylin sequence for nuclear visualization (5 min each), followed by tap water rinsing. Sections were then counterstained using a Ponceau–acid fuchsin solution for 5 min (prepared by mixing equal parts of Solution A and Solution B: Solution A, 0.5g Ponceau de xylidine in 1 mL glacial acetic acid and 100mL distilled water; Solution B, 0.5g acid fuchsin in 1mL glacial acetic acid and 100mL distilled water). After that, sections were differentiated in 1% phosphomolybdic acid for 4 min until collagen appeared colorless and counterstained with 1% light green SF for 2 min. Between each staining step, slides were rinsed with distilled water. Staining quality was microscopically confirmed at the differentiation and counterstaining stages.

Following all staining protocols, sections were dehydrated, cleared in xylene, mounted and cover slipped.

### Immunohistochemical Staining

Sections cut at 3 µm thickness were immunostained using either the Leica Bond III Autostainer (Leica Biosystems) or the Ventana Benchmark Ultra platform (Roche). Automated staining protocols were optimized to select the appropriate antigen retrieval method, antibody dilution and incubation period, and detection steps for each FRC antibody. Appropriate negative and positive control tissues were included with each batch.

PDGFRβ was stained using a recombinant rabbit monoclonal antibody (clone RM303) at 1:100 dilution, with heat-induced epitope retrieval (HIER) in pH 9 buffer for 30 min on the Leica Bond III platform.

Integrin β1 was detected using a rabbit monoclonal antibody (clone EP1041Y) at 1:300 dilution. Enzymatic antigen retrieval was performed using proteinase K for 10 min on the Leica Bond III platform.

FAP (clone EPR20021) staining was performed at 1:100 dilution. HIER in pH 9 buffer for 30 min on Leica Bond III platform was the most effective, but results were not consistent.

Podoplanin (clone D240) and SMA (clone 1A4) staining were performed using mouse monoclonal, ready to use, prediluted kits with HIER on the Ventana Benchmark Ultra platform.

### Region of Interest (ROI) Selection

Whole-slide images of all stained sections were acquired using the Glissando Desktop Scanner (Objective Imaging). Digital images were viewed using QuPath (0.5.1-x64).

For each PDGFRβ-stained lymph node section, ROIs measuring 250,000 µm² were selected at x20 magnification. In uninvolved (negative) nodes, eight ROI per section were manually selected from T cell zones, including a representative mix of paracortical and interfollicular areas, while avoiding germinal centers.

In metastatic (positive) lymph nodes, eight ROI were selected from areas of metastasis. Nodes containing only isolated tumor cells, micrometastases or small macrometastases were excluded, as the deposits were smaller than the standardised ROI area. For each positive node, an additional eight ROI were selected from residual lymphoid tissue, at least 500 µm away from metastatic tumor deposits. Nodes that were completely replaced by tumor were excluded from this residual tissue analysis.

In cases with histological evidence of NACT-induced fibrosis, four additional ROI were selected from fibrotic areas. Where fewer than the target number of ROI could be identified due to tissue limitations, the maximum feasible number was selected and documented.

### Image Analysis

All ROI were analyzed using the TWOMBLI plugin for ImageJ (NIH) ^27^ to quantify spatial metrics of the FRC network. Prior to analysis, the blue haematoxylin counterstain was digitally removed from PDGFRβ-stained ROI images to minimize background noise. The following optimized parameters were applied to all images: contrast saturation = 0.35, minimum line width = 5, maximum line width = 20, minimum branch length = 10 and maximum display HDMI = 237. TWOMBLI derived outputs were: lacunarity (a measure of how completely the FRC network fills the ROI), HDMI (proportion of the ROI covered by FRC network), HGU (the number of endpoints per unit length), fiber alignment, FRC fiber length, FRC fiber width, the number of FRC endpoints and the number of FRC branchpoints.

The same ROI images from metastatic ALN and areas of NACT-induced fibrosis were additionally analyzed using CurveAlign (v5.0, MATLAB-based), which is designed to accurately quantify fiber orientation and anisotropy.

Cell counts within each ROI were performed using QuPath, with automated detection parameters optimized for each staining condition.

## QUANTIFICATION AND STATISTICAL ANALYSIS

For each lymph node, the median value across the eight ROI was calculated for all quantitative parameters to minimize the impact of outliers. Data analysis was performed in RStudio (version 2024.09.1+394), using the following R packages: “tidyverse” for data manipulation and visualization, “ggplot2” for plotting, stats for regression analysis, and “survival” and “survminer” for survival analysis.

Associations between TWOMBLI derived matrix features and clinical variables were assessed using multivariate linear regression. For comparisons between unmatched, non-parametric categorical subgroups, the Kruskal–Wallis test was performed in GraphPad Prism (version 10.4.1). A p-value ≤ 0.05 was considered statistically significant.

The PCA included both TWOMBLI (i.e. lacunarity; branchpoints (normalized to fiber length); endpoints (normalized to fiber length); HDMI; and alignment) and clinical (i.e. NACT exposure; size/type of metastasis; pathological tumor stage; age; axillary tumor burden; and grade) parameters. Since some TWOMBLI parameters (i.e. HGU; fiber length; and width) were dependent on those listed above, they were excluded from the PCA.

Survival analysis was conducted in RStudio using the Kaplan–Meier method; group comparisons were performed using the Gehan-Breslow-Wilcoxon method.

