## Supplementary Data for "Topological Analysis of the Human Lymph Node Reticular Network Predicts Outcome in Breast Cancer"

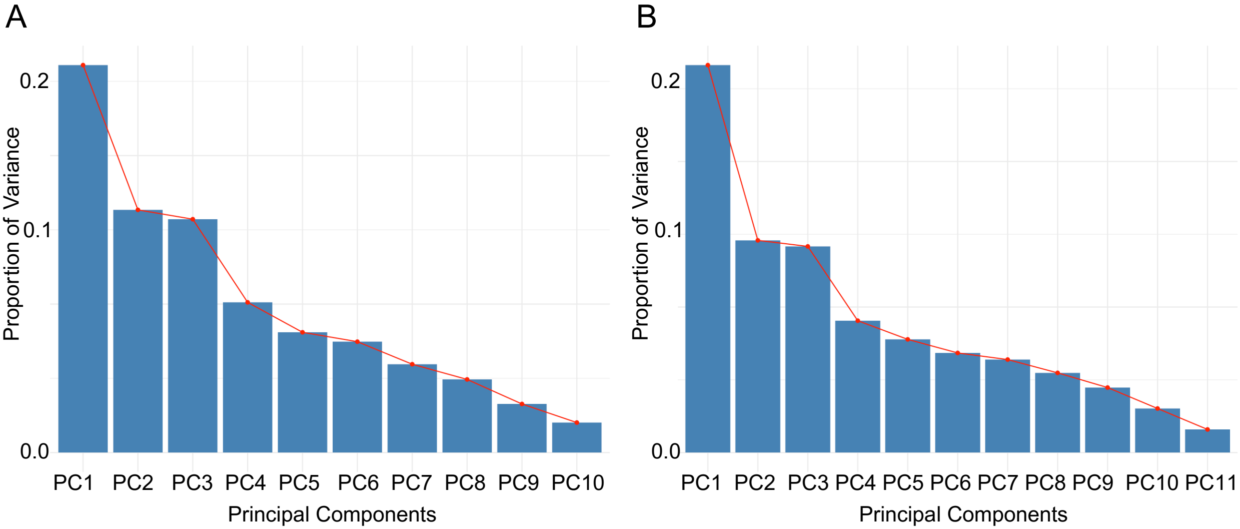
A
B
Proportion of Variance
0.2
0.1
0.0
PC1
PC2
PC3
PC4
PC5
PC6
PC7
PC8
PC9
PC10
PC11
Principal Components

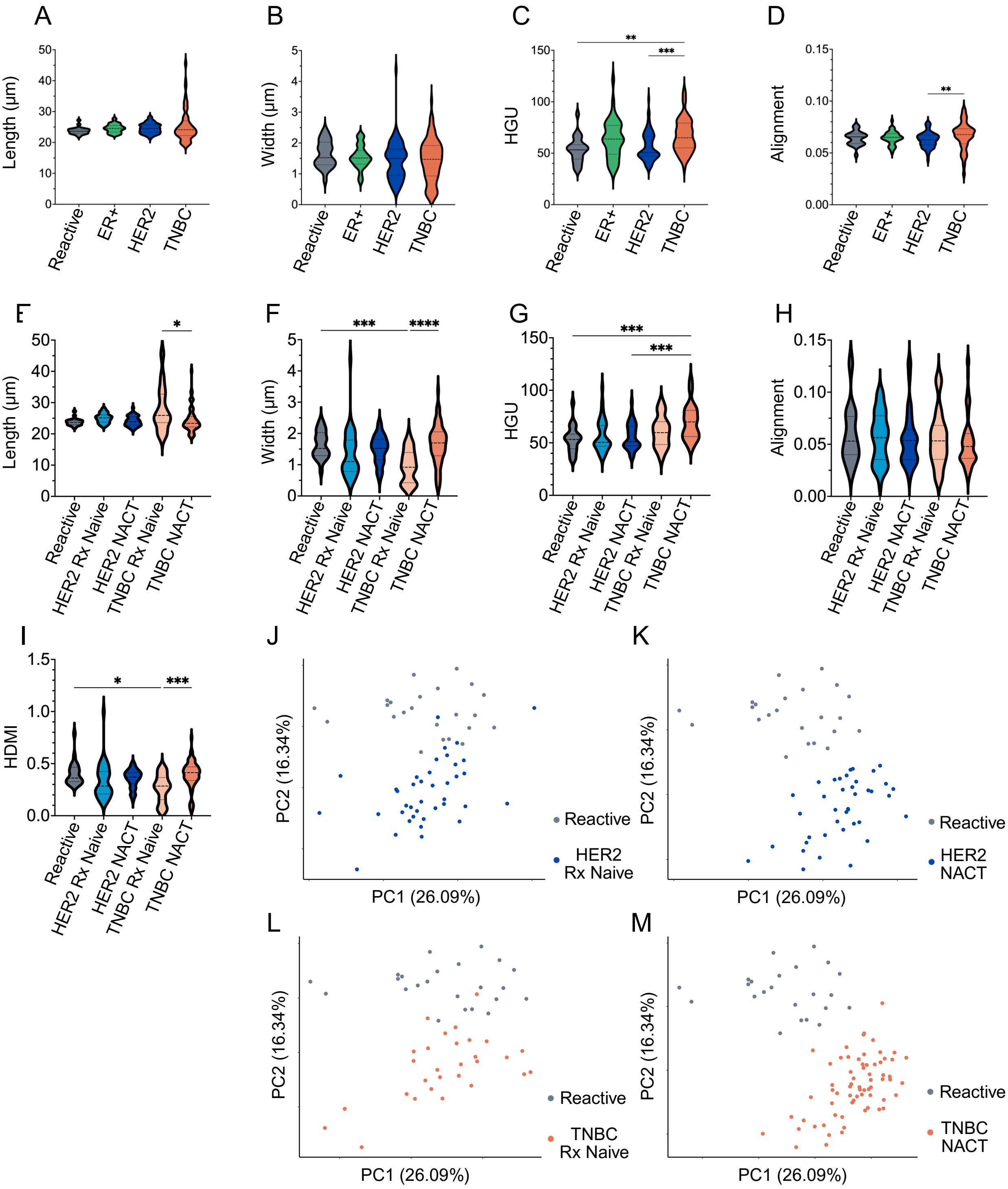
A
B
C
D
Length (μm)
Width (μm)
HGU
Alignment
Reactive
ER+
HER2
TNBC
E
F
G
H
HER2 Rx Naive
HER2 NACT
TNBC Rx Naive
TNBC NACT
I
HDMI
J
K
L
M
PC1 (26.09%)
PC2 (16.34%)
Reactive
HER2 Rx Naive
HER2 NACT
TNBC Rx Naive
TNBC NACT

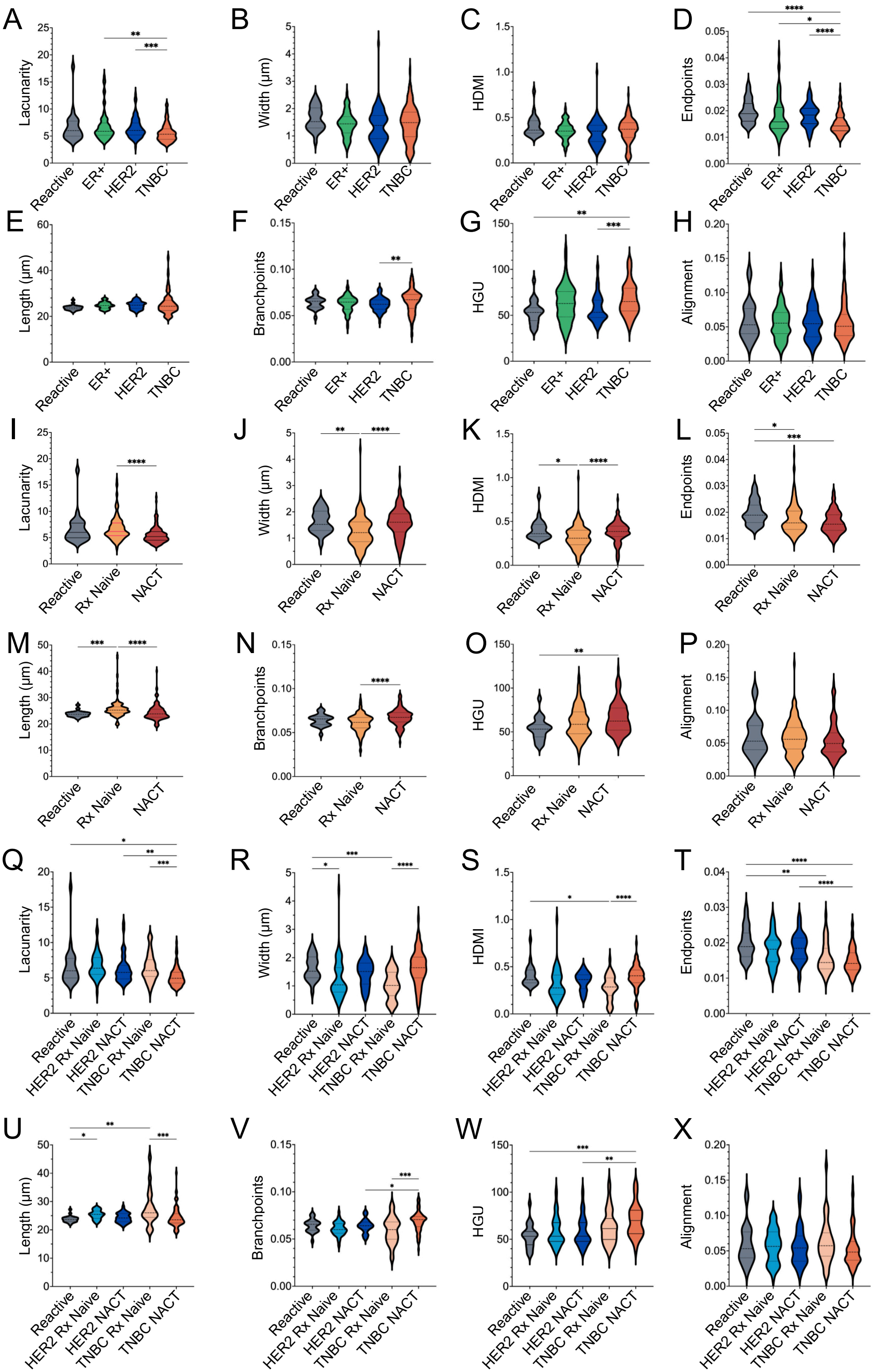
A
B
C
D
E
F
G
H
I
J
K
L
M
N
O
P
Q
R
S
T
U
V
W
X
Lacunarity
Width (µm)
HDMI
Endpoints
Length (µm)
Branchpoints
HGU
Alignment
Reactive
ER+
HER2
TNBC
Rx Naive
NACT
HER2 Rx Naive
HER2 NACT
TNBC Rx Naive
TNBC NACT

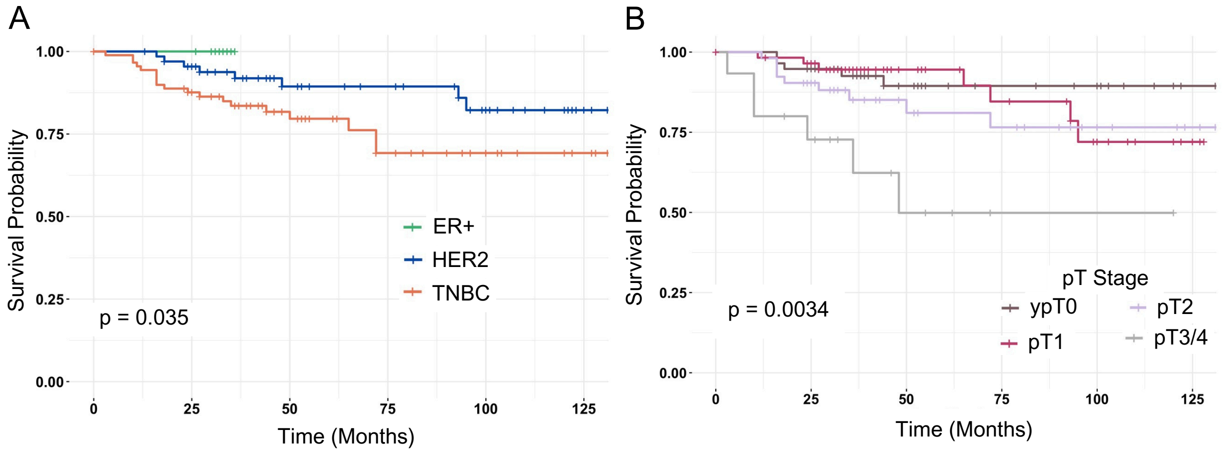

A
Survival Probability
1.00
0.75
0.50
0.25
0.00
0
25
50
75
100
125
Time (Months)
ER+
HER2
TNBC
p = 0.035
B
Survival Probability
Time (Months)
p = 0.0034
pT Stage
ypT0
pT1
pT2
pT3/4

### SUPPLEMENTARY FIGURE LEGENDS

#### Supplementary Figure 1. Differences Between Lymph Node Subsets Driven by Combined FRC Topological and Clinical Features

(A) Scree plot for PCA integrating TWOMBLI derived reticular network features and clinical variables from reactive, benign nodes and uninvolved lymph nodes, shown in **Figure 3**.

(B) Scree plot for PCA integrating TWOMBLI derived reticular network features and clinical variables from reactive, benign nodes, residual lymphoid tissue in involved nodes and uninvolved lymph nodes, shown in **Figure 4**.

#### Supplementary Figure 2. Stratified Analysis of FRC Network Remodelling in Uninvolved ALN by TNBC and NACT Status

(A-D) Violin plots illustrating molecular subtype-specific differences in uninvolved ALN FRC fiber length (μm) (A), width (μm) (B), HGU (C) and fiber alignment (D) (reactive: n = 23 ALN; ER+: n = 40 ALN; HER2: n = 78 ALN; TNBC: n = 96 ALN). Each data point represents the median of 8 ROI per node. **$p \leq 0.01$; ***$p \leq 0.001$; ****$p \leq 0.0001$ (Kruskal-Wallis test). Median with interquartile range, with minimum and maximum.

(E-I) Violin plots showing the impact of NACT on FRC fiber length (μm) (E), width (μm) (F), HGU (G), fiber alignment (H) and HDMI (I) in uninvolved ALN, stratified by molecular subtype of BC (reactive: n = 23 ALN; HER2 treatment-naïve: n = 39 ALN; HER2 post-NACT: n = 39 ALN; TNBC treatment-naïve: n = 29 ALN; TNBC post-NACT: n = 67 ALN). Each data point represents the median of 8 ROI per node. *$p \leq 0.05$; ***$p \leq 0.001$; ****$p \leq 0.0001$ (Kruskal-Wallis test). Median with interquartile range, with minimum and maximum.

(J and K) PCA comparing effects of NACT on uninvolved nodes from patients with HER2 BC (reactive: n = 23 ALN; treatment-naïve (J): n = 39 ALN; post-NACT (K): n = 39 ALN). Each data point represents the median of 8 ROI per node.

(L and M) PCA comparing effects of NACT on uninvolved nodes from patients with TNBC (reactive: n = 23 ALN; treatment-naïve (L): n = 29 ALN; post-NACT (M): n = 67 ALN). Each data point represents the median of 8 ROI per node.

#### Supplementary Figure 3. Stratified Analysis of FRC Network Remodelling in Uninvolved and Residual ALN

(A-X) Violin plots showing differences in TWOMBLI derived parameters of uninvolved and residual ALN FRC. For all plots, each data point represents the median of 8 ROIs per node. *$p \leq 0.05$; **$p \leq 0.01$; ***$p \leq 0.001$; ****$p \leq 0.0001$ (Kruskal-Wallis test). Median with interquartile range, with minimum and maximum.

(A-H) Violin plots showing molecular subtype-specific differences in FRC network lacunarity (A), fiber width (μm) (B), HDMI (C), number of endpoints (normalised to fiber length) (D), fiber length (μm) (E), number of branchpoints (normalised to fiber length) (F), HGU (G) and alignment (H) (reactive: n = 23 ALN; ER+: n = 69 ALN; HER2: n = 93 ALN; TNBC: n = 124 ALN).

(I-P) Violin plots showing effect of NACT on FRC network lacunarity (I), fiber width (μm) (J), HDMI (K), number of endpoints (normalised to fiber length) (L), fiber length (μm) (M), number of branchpoints (normalised to fiber length) (N), HGU (O) and alignment (P) (reactive: n = 23 ALN; treatment naïve: n = 150 ALN; post-NACT = 136 ALN).

(Q-X) Violin plots stratified by exposure to NACT and molecular subtype in FRC network lacunarity (Q), fiber width (μm) (R), HDMI (S), number of endpoints (normalised to fiber length) (T), fiber length (μm) (U), number of branchpoints (normalised to fiber length) (V), HGU (W) and alignment (X) (reactive: n = 23 ALN; HER2 treatment naïve: n = 49 ALN; HER2 post-NACT: n = 44 ALN; TNBC treatment naïve: n = 41 ALN; TNBC post-NACT: n = 83 ALN).

**Supplementary Figure 4. Clinical Stratification Confirms Prognostic Impact of Molecular Subtype and Tumor Stage**

(A) Kaplan–Meier survival analysis, stratified by molecular subtype demonstrating that patients with TNBC had the worst prognosis ($p = 0.035$; ER+: $n = 25$ patients; HER2: $n = 67$ patients; TNBC: $n = 87$ patients; Gehan-Breslow-Wilcoxon method).

(B) Kaplan–Meier survival analysis stratified by pathological tumor stage (pT), showing that patients with tumors of a higher stage had worse prognosis ($p = 0.0034$; ypT0 (pathological tumor stage 0 after neoadjuvant chemotherapy): $n = 55$ patients; pT1: $n = 59$ patients; pT2: $n = 52$ patients; pT3/4: $n = 13$ patients; Gehan-Breslow-Wilcoxon method).

### SUPPLEMENTARY TABLES

#### Supplementary Table 1. Clinico-pathological Characteristics of Reactive Patient Cohort

| Clinicopathological Characteristic | No. (%) |
|---|---|
| **Age** | |
| 30 - 40 | 3 (13) |
| 40 - 50 | 2 (9) |
| 50 - 60 | 3 (13) |
| 60 - 70 | 4 (17) |
| 70 - 80 | 2 (9) |
| 80 - 90 | 9 (39) |
| **Site of Node** | |
| Axilla | 11 (48) |
| Cervical | 6 (26) |
| Jugular | 4 (17) |
| Upper limb (not further specified) | 2 (9) |
| **Type of Biopsy** | |
| Core biopsy | 12 (52) |
| Excisional biopsy | 11 (48) |
| **Diagnosis** | |
| Reactive lymphoid hyperplasia | 9 (39) |
| Reactive follicular hyperplasia | 8 (35) |
| Reactive lymph node (not further specified) | 4 (17) |
| Dermatopathic lymphadenopathy | 2 (9) |

#### Supplementary Table 2. Summary of Linear Regression Model Showing Predictive Power of Age in Reactive Cohort

| TWOMBLI output | R Score | F Statistic | P-value |
|---|---|---|---|
| **Branchpoints** | 0.10 | 2.39 | 0.137 |
| **Endpoints** | 0.15 | 3.47 | 0.076 |
| **Lacunarity** | 0.14 | 3.44 | 0.078 |
| **Alignment** | 0.03 | 0.76 | 0.394 |
| **Fibre Width** | 0.06 | 1.36 | 0.257 |
| **Fibre Length** | 0.06 | 1.33 | 0.262 |
| **HGU** | 0.01 | 0.22 | 0.647 |
| **HDMI** | 4.361e-06 | 9.159e-05 | 0.993 |

Abbreviations: High density matrix intensity (HDMI), hyphal growth units (HGU)

**Supplementary Table 3. Results of Linear Multi-Variate Analysis for Uninvolved Nodes**

| TWOMBLI output | R Score | F Statistic | P-value | Significant Predictors (p-value) |
|---|---|---|---|---|
| **Endpoints (normalised)** | 0.127 | 6.748 | 0.000007 | Molecular Subtype (0.0006), Tumour burden in axilla (0.01) |
| **Fibre Width** | 0.114 | 5.961 | 0.00003 | NACT (0.00006) |
| **Branchpoints (normalised)** | 0.109 | 5.656 | 0.00006 | NACT (0.00009), Grade (0.04) |
| **Lacunarity** | 0.105 | 5.468 | 0.00009 | NACT (0.002), Molecular subtype (0.02) |
| **Fibre Length** | 0.090 | 4.565 | 0.0005 | NACT (0.0007), Tumour burden in axilla (0.02) |
| **HDMI** | 0.088 | 4.474 | 0.0007 | NACT (0.001) |
| **HGU** | 0.073 | 3.560 | 0.004 | NACT (0.05), Tumour burden in axilla (0.04) |
| **Alignment** | 0.011 | 0.517 | ns | None |

Abbreviations: High density matrix intensity (HDMI), hyphal growth units (HGU), neoadjuvant chemotherapy (NACT).

**Supplementary Table 4. PCA Loadings for Uninvolved Nodes**

| Variable | PC1 Loading | PC2 Loading | PC3 Loading |
|---|---|---|---|
| **Lacunarity** | -0.506 | -0.129 | 0.312 |
| **Branchpoints (normalised)** | 0.478 | 0.187 | -0.219 |
| **NACT** | 0.403 | -0.271 | 0.329 |
| **Endpoints (normalised)** | -0.276 | 0.285 | 0.323 |
| **Age** | -0.265 | 0.410 | -0.288 |
| **HDMI** | 0.244 | 0.343 | 0.003 |
| **Alignment** | -0.241 | -0.301 | 0.149 |
| **pT** | -0.189 | -0.147 | -0.556 |
| **Tumour burden in axilla** | -0.179 | -0.239 | -0.470 |
| **Grade** | 0.146 | -0.582 | -0.079 |

Abbreviations: High density matrix intensity (HDMI), neoadjuvant chemotherapy (NACT), pathological tumour stage (pT).

### Supplementary Table 5. Results of Linear Multi-Variate Analysis for Residual and Uninvolved Nodes

| TWOMBLI output | R Score | F Statistic | P-value | Significant Predictors (p-value) |
|---|---|---|---|---|
| **Lacunarity** | 0.157 | 9.343 | 0.000000002 | Size/type of met (0.00001), NACT (0.001), Molecular subtype (0.008) |
| **Fibre Width** | 0.125 | 7.179 | 0.0000004 | NACT (0.00002), Size/type of met (0.04) |
| **HDMI** | 0.122 | 5.210 | 0.00004 | NACT (0.001) |
| **Endpoints (normalised)** | 0.112 | 6.374 | 0.0000002 | Molecular subtype (0.0006), Tumour burden in axilla (0.004) |
| **Fibre Length** | 0.092 | 5.087 | 0.00005 | NACT (0.00009), Tumour burden in axilla (0.02), Molecular subtype (0.04) |
| **Branchpoints (normalised)** | 0.010 | 6.862 | 0.0000008 | NACT (0.00002), Size/type of met (0.05) |
| **HGU** | 0.072 | 3.895 | 0.0009 | Tumour burden in axilla (0.01). NACT (0.02) |
| **Alignment** | 0.026 | 1.127 | ns | Size/type of met (0.03) |

Abbreviations: High density matrix intensity (HDMI), neoadjuvant chemotherapy (NACT), pathological tumour stage (pT).

### Supplementary Table 6. PCA Loadings for Residual and Uninvolved Nodes

| Variable | PC1 Loading | PC2 Loading | PC3 Loading |
|---|---|---|---|
| **Lacunarity** | 0.410 | 0.256 | -0.124 |
| **Branchpoints (normalised)** | -0.467 | -0.182 | 0.154 |
| **Endpoints (normalised)** | 0.336 | 0.416 | 0.104 |
| **NACT** | -0.336 | 0.218 | -0.421 |
| **HDMI** | -0.312 | 0.024 | 0.264 |
| **Size/type of met** | 0.234 | -0.286 | -0.161 |
| **pT** | 0.219 | -0.516 | -0.014 |
| **Alignment** | 0.208 | 0.025 | -0.234 |
| **Age** | 0.176 | -0.162 | 0.542 |
| **Tumour burden in axilla** | 0.162 | -0.516 | -0.062 |
| **Grade** | -0.077 | -0.194 | -0.569 |

Abbreviations: High density matrix intensity (HDMI), neoadjuvant chemotherapy (NACT), pathological tumour stage (pT).
